# Resistance to tembotrione, a 4- Hydroxyphenylpyruvate Dioxygenase-Inhibitor in *Sorghum bicolor*

**DOI:** 10.1101/2020.07.29.227512

**Authors:** Balaji Aravindhan Pandian, Aruna Varanasi, Amaranatha R. Vennapusa, Rajendran Sathishraj, Guifang Lin, Mingxia Zhao, Madison Tunnell, Tesfaye Tesso, Sanzhen Liu, P.V. Vara Prasad, Mithila Jugulam

## Abstract

Postemergence grass weed control continues to be a big challenge in grain sorghum (*Sorghum bicolor* L. Moench), primarily due to a lack of herbicide options registered for use in this crop. The development of herbicide-resistant sorghum technology to facilitate broad-spectrum postemergence weed control is an economical and viable solution. The 4-hydroxyphenylpyruvate dioxygenase-inhibitor herbicides (e.g. mesotrione or tembotrione) can control broad-spectrum of weeds including grasses, which, however, is not registered for postemergence application in sorghum due to crop injury. In this study we identified two tembotrione-resistant sorghum genotypes (G-200, G-350) and one highly susceptible genotype (S-1) through screening 317 sorghum lines from the sorghum association panel (SAP). Compared to S-1, G-200 and G-350 exhibited 10- and 7-fold more resistance to tembotrione, respectively. Genetic analyses of the F_1_ and F_2_ progeny generated from a cross between tembotrione-resistant and -susceptible genotypes demonstrated that the resistance is a semi-dominant polygenic trait. Furthermore, cytochrome P450 (CYP)-inhibitor assay using malathion and piperonyl butoxide suggested possible CYP-mediated metabolism of tembotrione in G-200 and G-350. Genotype-by-sequencing based quantitative trait loci (QTL) mapping revealed eight QTLs associated with tembotrione resistance in grain sorghum. Sorghum genotypes G-200 and G-350 confer a high level of metabolic resistance to tembotrione and controlled by a polygenic trait. There is an enormous potential to introgress the tembotrione resistance into breeding lines to develop agronomically desirable sorghum hybrids.

**One-sentence summary:** This research focuses on characterization, genetic analyses, identification of QTLs-linked to metabolic resistance to tembotrione in Sorghum bicolor, for improved weed control and increased yield

## Introduction

Grain sorghum [*Sorghum bicolor* (L.) Moench ssp. *bicolor*] is one of the most versatile crops with multiple uses including for food, feed, and fuel (Ciampitti and Prasad, 2019). Sorghum performs better than corn (*Zea mays*) under rainfed and low input conditions (Valadabad *et al.*, 2000; Staggenborg *et al.*, 2008). The US is the largest producer of grain sorghum; and almost half of the US grain sorghum is produced in Kansas (USDA-NASS, 2020). Sorghum is primarily grown for cattle feed and ethanol production in the US, whereas, it is a staple food for millions of people in Africa, India, and South America (Taylor *et al.*, 2006; Dahlberg *et al.*, 2012). Weed infestation, specifically grass weed species pose a major problem in sorghum production and can reduce the crop yields up to 60%, if left uncontrolled (Thompson *et al.*, 2019; Dille *et al.*, 2020). Palmer amaranth (*Amaranthus palmeri*), common waterhemp (*Amaranthus tuberculatus*), kochia (*Bassia scoparia*), common ragweed (*Ambrosia artemisiifolia*), common lambsquarters (*Chenopodium album*) are the major broadleaf weeds and johnsongrass (*Sorghum halepense*), shattercane (*Sorghum bicolor* ssp. *verticilliflorum*), large crabgrass (*Digitaria sanguinalis*) are major grass weeds found in grain sorghum fields (Stahlman *et al.*, 2000; Smith *et al.*, 2010). A wide range of postemergence (POST) herbicides are available to control broad-leaved weeds in sorghum. However, herbicide options for POST control of grasses are limited due to the susceptibility of sorghum to commonly used grass control herbicides (Thompson *et al.*, 2019).

The 4-hydroxyphenylpyruvate dioxygenase (HPPD)-inhibitors (e.g. mesotrione or tembotrione) are widely used to control broad-spectrum of weeds including grasses in corn because it can effectively metabolize HPPD-inhibitors (Williams and Pataky, 2010). However, these herbicides are not registered as POST in sorghum due to crop injury. Although these herbicides are widely used, till date only two weed species i.e., Palmer amaranth and common waterhemp have been documented to have evolved resistance to HPPD-inhibitors (Heap, 2020). These herbicides inhibit the HPPD enzyme, which is important for the conversion of 4-hydroxyphenyl pyruvate to homogentisate, an intermediate in plastoquinone and tocopherol biosynthesis pathway in plants (Lee *et al.*, 1998). Plastoquinone is essential for the carotenoid biosynthesis, which protects the chlorophyll by absorbing excited electrons released during photosynthesis. Depletion of carotenoids causes damage to the chlorophyll by photo-oxidation resulting in bleaching followed by necrosis and plant death (Dankov *et al.*, 2009). HPPD-inhibitors include four chemical families isoxazole, pyrazole, pyrazolone, and triketones, and were introduced in the 1980s for weed control (van Almsick, 2009).

Herbicide resistance in plants can be conferred by two major mechanisms: a) target-site resistance (TSR): mutation(s) in the herbicide target gene leading to the reduced affinity of the target enzyme for herbicide binding or due to increased expression of target enzyme; and b) non-target site resistance (NTSR): increased metabolism or reduced absorption/translocation of herbicides (Gaines *et al.*, 2020). Metabolism of HPPD-inhibitors by cytochrome P450 enzyme (CYPs) activity is the most common mechanism of resistance found in crops as well as weeds (Ahrens *et al.*, 2013). Nonetheless, increased expression of *HPPD*-gene has also been reported in some biotypes of Palmer amaranth (Nakka *et al.*, 2017). Recently, a modified *HPPD-*gene from *Pseudomonas fluorescens* and *Avena sativa* which is insensitive to HPPD-inhibitors was used to develop transgenic soybeans (*Glycine max*) resistant to HPPD-inhibitors by Bayer Crop Science (Matringe *et al.*, 2005; Dreesen *et al.*, 2018) and Syngenta (Hawkes *et al.*, 2016), respectively. Dupont-Pioneer used an insensitive shuffled variant of corn *HPPD*-gene that confers a high level of resistance to HPPD-inhibitors in soybean (Siehl *et al.*, 2014).

CYPs are one of the largest enzyme families involved in xenobiotic metabolism in microorganisms, insects, plants, and humans imparting resistance, respectively, to antibiotics, insecticide, and herbicide, and drugs (Pandian *et al.*, 2020; Siminszky, 2006). The activity of CYPs can be inhibited using several chemical compounds; 1-aminobenzo-triazole (ABT), tetcyclacis (TET), piperonyl butoxide (PBO), tridiphane, and organophosphate insecticides such as malathion, phorate (Siminszky, 2006; Busi *et al.*, 2017). Treatment with CYP-inhibitors before herbicide application will competitively reduce the CYP activity resulting in decreased metabolism of herbicide, thereby, reducing the level of resistance (Siminszky, 2006). CYP-inhibitors have been widely used to determine metabolic resistance to herbicides in several plant species. Specifically, malathion and PBO were used to demonstrate the inhibition of CYP activity and the reversal of crop tolerance to HPPD-inhibitors in corn (Ma *et al.*, 2013; Oliveira *et al.*, 2018).

Development of sorghum hybrids resistant to HPPD-inhibitors will provide POST herbicide option to control grass weeds (Thompson *et al.*, 2019). Tembotrione is a triketone herbicide which has broad spectrum activity including grass weeds. Furthermore, the efficacy of tembotrione is high on grass weeds compared to other triketones (Ahrens *et al.*, 2013). Mesotrione, a triketone herbicide similar to tembotrione is registered for preemergence (PRE) use in sorghum but not as POST; however, tembotrione is not registered for PRE or POST usage in sorghum. We hypothesize that screening diverse genotypes from the sorghum association panel (SAP) will facilitate identification of genotypes resistant to tembotrione; and such resistance, similar to maize, is associated with CYP-mediated metabolism. The specific objectives of this research were to identify and characterize sorghum genotypes with resistance to tembotrione, to investigate the inheritance and mechanism of resistance to tembotrione, and to identify genetic loci conferring tembotrione resistance.

## Results

### Identification of tembotrione resistant genotypes

Among 317 genotypes from the SAP initially screened under *in vitro* (tissue culture) conditions, 10 genotypes showed ≤ 70% tembotrione injury at 2 weeks after treatment (WAT) with significant recovery by 4 WAT (Table S1). One genotype, S-1 was found highly sensitive to tembotrione as these plants did not survive tembotrione treatment. In response to the tembotrione application at 92 g ai ha^−1^ (field recommended dose of tembotrione), under greenhouse conditions, out of the above10 genotypes, only two, i.e., G-200 and G-350 showed the least injury at 2 WAT (73 and 70%, respectively) and the smallest dry biomass reduction at 3 WAT (25.36 and 25.77%, respectively) (Table S2). In the treated plants, the tembotrione injury symptoms appeared by 2 WAT, after which in G-200 and G-350 the symptoms dissipated and plants started to recover. However, leaf chlorosis and bleaching symptoms, typical of tembotrione injury, were visible on all 10 genotypes at variable degree. The sensitive genotype, S-1, and Pioneer 84G62 (a commercially used sorghum hybrid) exhibited 100 and 90% injury with a biomass reduction of 2.69 and 11.63%, respectively (Table S2).

### Tembotrione dose-response assay

Based on the GR_50_ (dose required for 50% growth reduction) values the resistant genotypes, G-200 and G-350 required 206 and 104 g ai ha^−1^ of tembotrione, respectively, for 50% growth reduction which is higher than the field recommended dose of 92 g ai ha^−1^; whereas, Pioneer 84G62 and S-1 required only 24 and 38 g ai ha^−1^ for the same level of growth reduction. The GR_50_ values of G-200, G-350, and Pioneer 84G62 were significantly different from S-1. On the basis of GR_50_ values, compared to S-1, the genotypes G-200, G-350, and Pioneer 84G62 were ~10x, 7x, and 1.5x more resistant, respectively, to tembotrione (Figure 1; Table 1).

**Table 1.** Regression parameters describing the response of sorghum genotypes to tembotrione under greenhouse and field conditions and (GR_50_: dose required for 50% growth reduction; ID_50_ dose required for 50% visual injury; SE: standard error; RI: Resistance index R/S; Resistant/Susceptible).

| Genotype | Greenhouse |  | Field |  |
| --- | --- | --- | --- | --- |
|  | GR<sub>50</sub> (SE) | RI (R/S) | ID<sub>50</sub> (SE) | RI (R/S) |
| S-1 | 22.1 (3.4) | 1 | 68.4 (14.3) | 1 |
| Pioneer 84G62 | 36.7 (5.6) | 2 | 62.4 (51.0) | 1 |
| G-350 | 154.4 (68.8) ** | 7 | 96.6 (17.8) * | 1.5 |
| G-200 | 215.2 (160) * | 10 | 129.6 (71.8) ** | 2 |

Significantly different from S-1 at \*p ≤ 0.05 \*\*p ≤ 0.01

**Figure 1:**
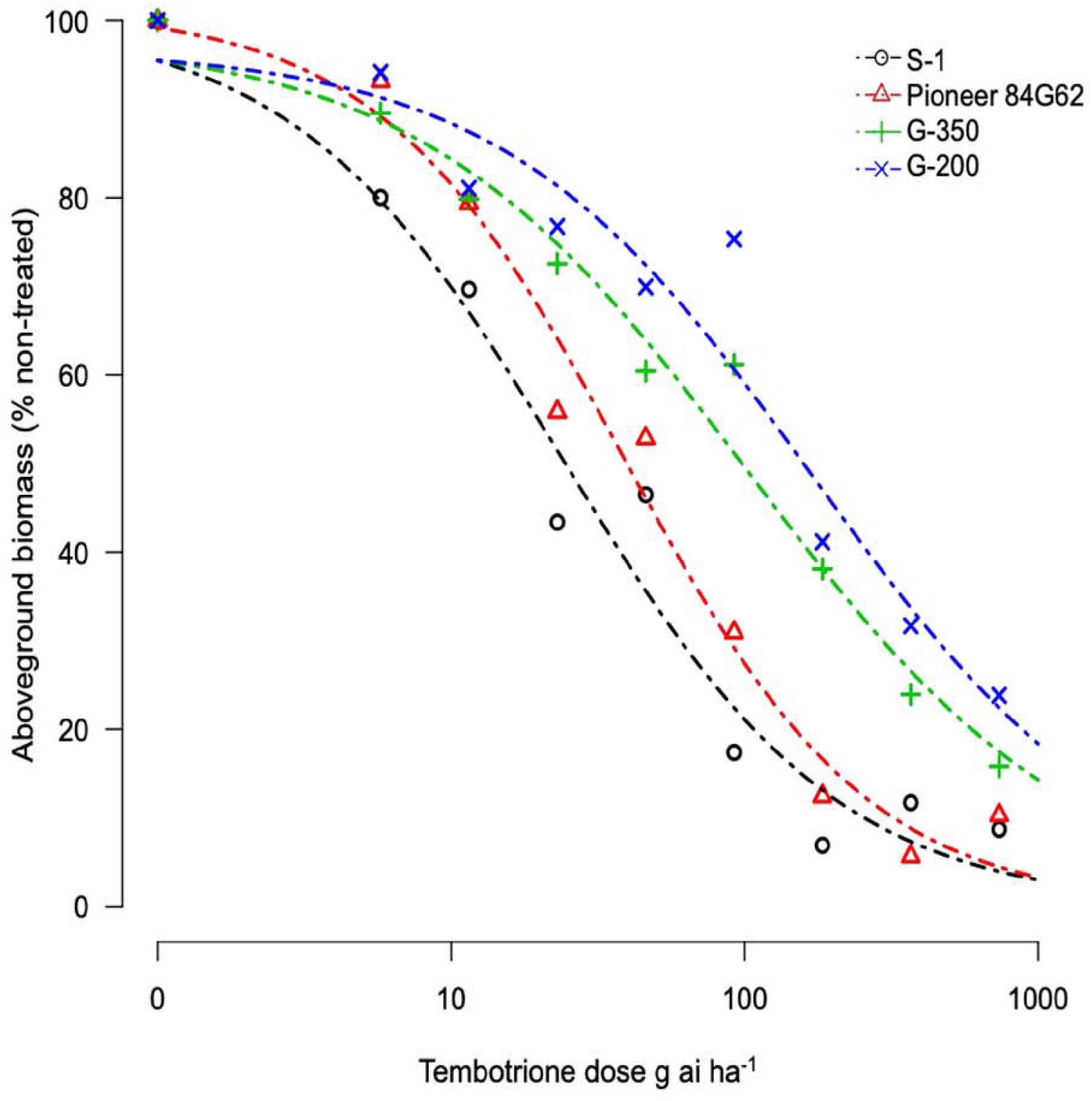
Tembortrine dose-response curves obtained by non-linear regression analysis of above-ground by biomass of S-1 (susceptible), Pioneer 84G6S (commercial hybrid), G-200, and G-350 (resistant) using the three-parameter log-ligistic model.

### Field testing

The response of sorghum genotypes, G-200 and G-350 along with S-1 and Pioneer 84G62 to POST application of tembotrione was tested under field conditions at two sites, Manhattan and Hays, KS. Since site by herbicide dose interaction was non-significant, data for tembotrione injury and yield were pooled and averaged across the two sites. In response to the tembotrione (POST) application, sorghum genotypes showed leaf chlorosis and bleaching symptoms followed by necrotic lesions in sensitive plants. The tembotrione injury was visible on all 4 sorghum genotypes at 1 WAT. A dose-response curve was generated using the injury ratings at 2 WAT. On the basis of percent injury at 2 WAT and ID_50_ (dose required for 50% visual injury) values, the genotype G-200 required the highest dose (129 g ai ha^−1^) of tembotrione followed by G-350 (96 g ai ha^−1^) genotype. Susceptible genotypes S-1 (62 g ai ha^−1^) and Pioneer 84G62 required the lowest dose of tembotrione (68 g ai ha^−1^) that caused 50% injury (Figure 2; Table 1). Days required for 50% recovery, ED_50_ from herbicide damage was also calculated. The genotypes, G-200 and G-350, recovered with no injury symptoms by 6 WAT at all doses of tembotrione, whereas, Pioneer 84G62 and S-1 took 8 weeks for recovery from 1x and 2x of application dose, and S-1 did not survive 4x dose of tembotrione (Table 2). In response to 1x dose of tembotrione, no significant reduction in grain yield of G-200 and G-350 was found; whereas, the susceptible genotypes, S-1 and Pioneer 84G62 showed ~40 and 30% grain yield reduction (Table 2) compared to non-treated. However, when treated with 2x- and 4x-doses of tembotrione, grain yields were significantly reduced in all genotypes (Table 2).

**Table 2.** Regression parameters describing the response of sorghum genotypes and their F1 progeny to tembotrione under greenhouse conditions (GR_50_: dose required for 50% growth reduction; SE: standard error; RI: Resistance index R/S; Resistant/Susceptible).

| Genotype | GR<sub>50</sub> (SE) | RI (R/S) |
| --- | --- | --- |
| S-1 | 36 (4) | 1 |
| S-1xG-200 | 104 (27) ** | 2.8 |
| S-1xG-350 | 117 (25) ** | 3.2 |
| G-350 | 172 (33) ** | 4 |
| G-200 | 218 (60) ** | 6 |

Significantly different from S-1 at \* $p \leq 0.05$ \*\* $p \leq 0.01$

**Figure 2:**
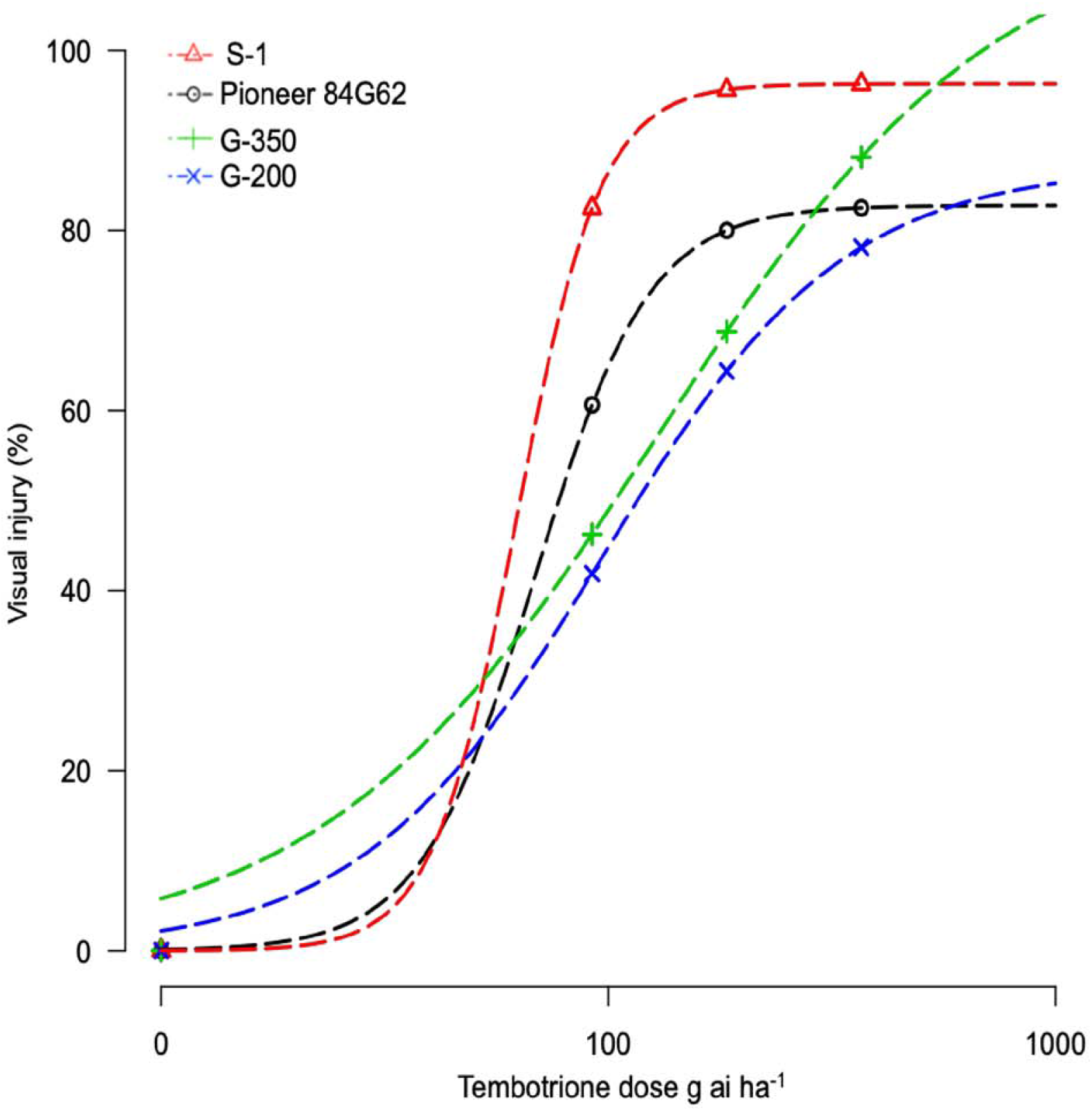
Tembortrine dose-response curves representing the percent injury of S-1 (susceptible), Pioneer 84G6S (commercial hybrid), G-200, and G-350 (resistant genotypes) in response to different doses of tembotrione at 2 weeks after treatment under field conditions.

### Mechanism of tembotrione resistance

Upon sequencing the *HPPD* gene from G-200, G-350, and S-1, no mutations were identified in the coding region of the *HPPD* gene (Data S1), suggesting that no target site alterations confer resistance to tembotrione in G-200 or G-350. In response to malathion or PBO followed by tembotrione treatments, G-200 and G-350 exhibited significant biomass reduction compared to plants treated with tembotrione alone. Malathion or PBO without tembotrione treatment had no effect on the sorghum genotypes tested (Figures 3 and 4). The corn inbred line B73 (known to be resistant to tembotrione) did not show any significant biomass reduction at 92, or 184 g ai ha^−1^ of tembotrione application (Figure 3a); however, exhibited a significant reduction in biomass in response to pre-treatment with malathion followed by 184 and 368 g ai ha^−1^ of tembotrione (Figure 3b). The genotype G-200 treated with malathion followed by 92, 184, and 368 g ai ha^−1^ showed more than 50% reduction in biomass compared to plants treated only with tembotrione. There is no significant difference in biomass accumulation when treated with 2000 or 4000 g ai ha^−1^ of malathion, except in malathion followed by 368 g ai ha^−1^ tembotrione treatment (Figure 3B). Whereas, G-350 showed significant biomass reduction only when treated with malathion followed by 92 g ai ha^−1^ tembotrione. The susceptible genotype, S-1, showed significant growth reduction at all doses of tembotrione or when pre-treated with malathion (Figure 3b). The corn B73 exhibited significant biomass reduction in response to PBO followed by 92, 184, and 368 g ai ha^−1^ of tembotrione treatment (Figures 4a and 4b). Both G-200 and G-350 genotypes also showed a significant reduction in biomass when pretreated with PBO followed by all doses of tembotrione. The S-1 was sensitive to all treatments applied (Figure 4b).

**Figure 3:**
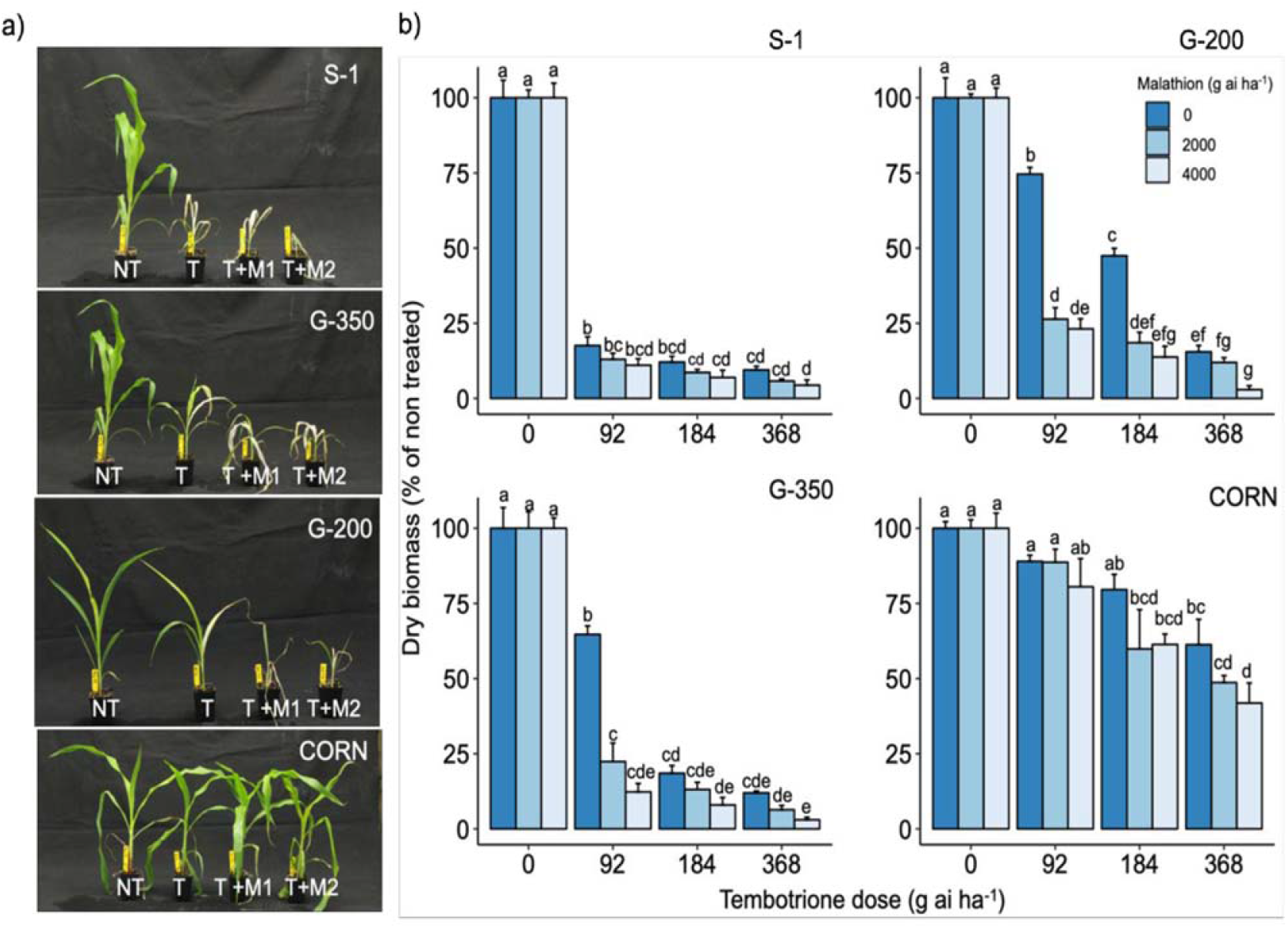
a) The response of S-1 (susceptible), G-200, G-350 (resistant) and, corn to field recommended dose tembotrione (92 g ai ha^−1^)(T) or malathion treatment (M1: 2000 g ai ha^−1^; M2: 4000 g ai ha^−1^) followed by field recommended dose tembotrione; NT: Non-treated; and b) Aboveground dry biomass of sorghum genotypes when pre-treated with malathion or different doses of tembotrione. The error bars represent the standard error; different alphabets indicate a significant difference between treatments (p ≤ 0.05).

**Figure 4:**
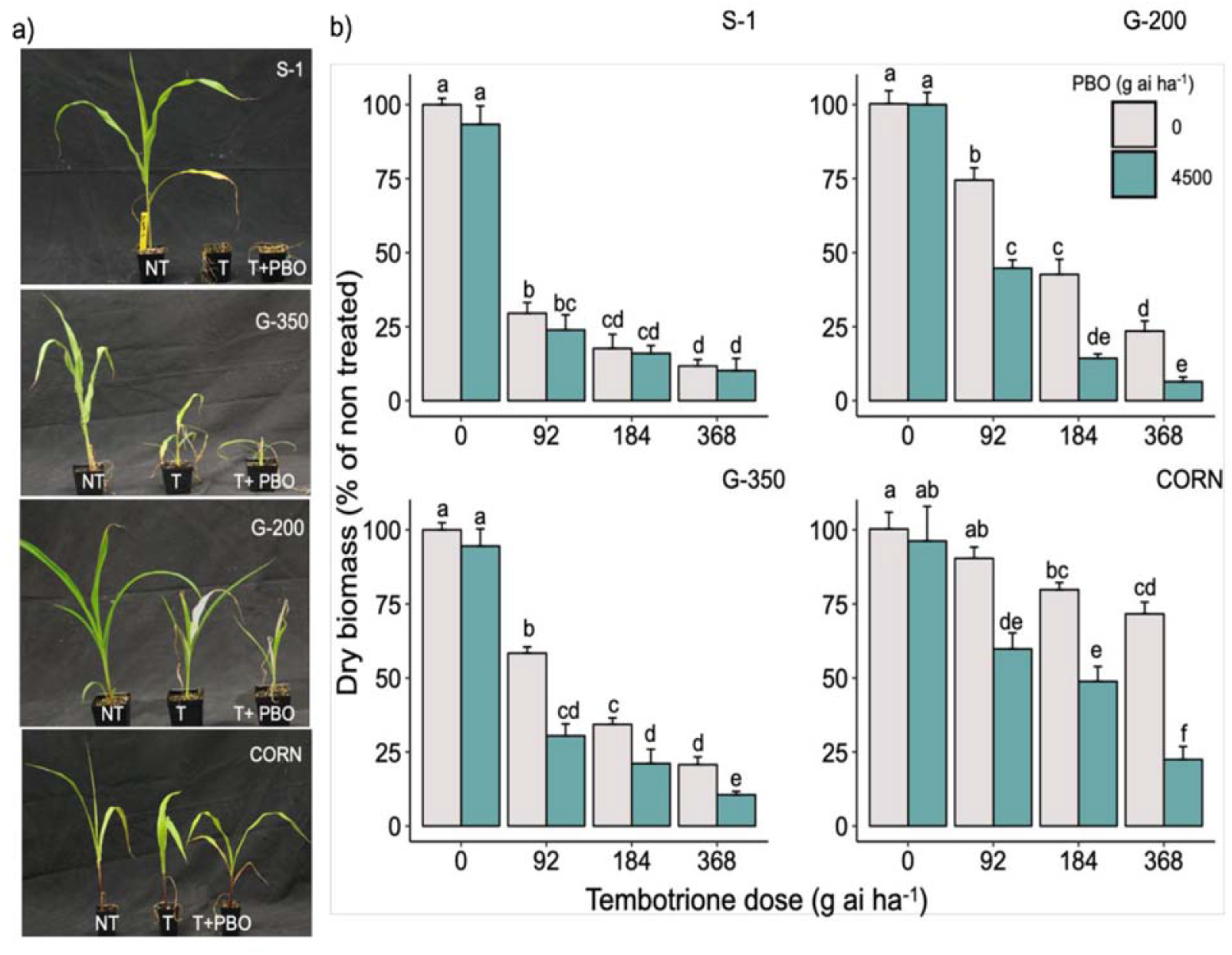
The response of S-1 (susceptible), G-200, G-350 (resistant) and, corn to field recommended dose tembotrione (92 g ai ha^−1^)(T) or Piperonyl butoxide (PBO) treatment (M1: 4500 g ai ha^−1^; followed by field recommended dose tembotrione; NT: Non-treated; and b) Aboveground dry biomass of sorghum genotypes when pre-treated with PBO or different doses of tembotrione. The error bars represent the standard error (n=8); different alphabets indicate a significant difference between treatments (p ≤ 0.05).

### Inheritance of tembotrione resistance

The F_1_ progeny of S-1 × G-200 and S-1 × G-350 showed an intermediate response relative to parents when treated with several doses of tembotrione. The GR_50_ of S-1 × G-200 and S-1 × G-350 were estimated at 104 and 117 g ai ha^−1^, respectively, which were less than their respective tembotrione-resistant parents, i.e., G-200 (218 g ai ha^−1^) and G-350 (172 g ai ha^−1^) (Figures 5 and 6; Table 3), suggesting that tembotrione resistance is a semi-dominant trait. The F_2_ progeny exhibited a continuous variation for tembotrione injury and recovery. Therefore, to perform a chi-square test frequency of segregation of tembotrione resistance or susceptibility in F_2_ progeny, the plants that had more than 80% tembotrione injury were grouped as susceptible and others as resistant. The observed segregation of resistant: susceptible (R:S) ratios from both the crosses did not comply with the expected ratios of 3:1 (R:S) for a single gene inherited trait, indicating that more than one gene is involved in tembotrione resistance in G-200 or G-350 genotypes of sorghum (Table 4).

**Table 3.** Recovery (ED_50_) and single plant yield (% of non-treated) in response to different doses of tembotrione under field conditions. (ED_50_: days required for 50% recovery; SE: standard error.

| Herbicide<br>Dose<br>(g ai ha<sup>-1</sup>) | ED<sub>50</sub>(SE) |  |  |  | Yield (%) |  |  |  |
| --- | --- | --- | --- | --- | --- | --- | --- | --- |
|  | S-1 | Pioneer<br>84G62 | G-350 | G-200 | S-1 | Pioneer<br>84G62 | G-200 | G-350 |
| 0 | - | - | - | - | 100a | 100a | 100a | 100a |
| 92 | 28 (6) | 23 (2) | 22(3) * | 18 (1) * | 53.7de | 68.5cd | 95.3ab | 99.8a |
| 184 | 30 (10) | 25 (3) | 21 (2) * | 20 (1) * | 19.4g | 45.4cd | 72.2cd | 78.0bc |
| 368 | - | 27 (1) | 21 (3) | 19 (1) | - | 14.6g | 25.3fg | 33.3fg |

Significantly different from S-1 at \* $p \leq 0.05$ . Numbers with different letters are significantly different; and numbers with same letters are not significantly different.

**Table 4.** Chi-Square analysis of tembotrione resistance at four weeks after treatment (R: Resistant and S: Susceptible).

| Cross | Run | Total | R | S | P-value |
| --- | --- | --- | --- | --- | --- |
| S-1 x G-200 | 1 | 200 | 168 | 32 | 0.00329** |
|  | 2 | 150 | 124 | 26 | 0.0372* |
|  | Run 1 and 2<br>Combined | 350 | 292 | 58 | 0.00033** |
| S-1 x G-350 | 1 | 220 | 197 | 23 | 0.00001** |

Significantly different at \*\* $\leq 0.01$ \* $p \leq 0.05$

**Figure 5:**
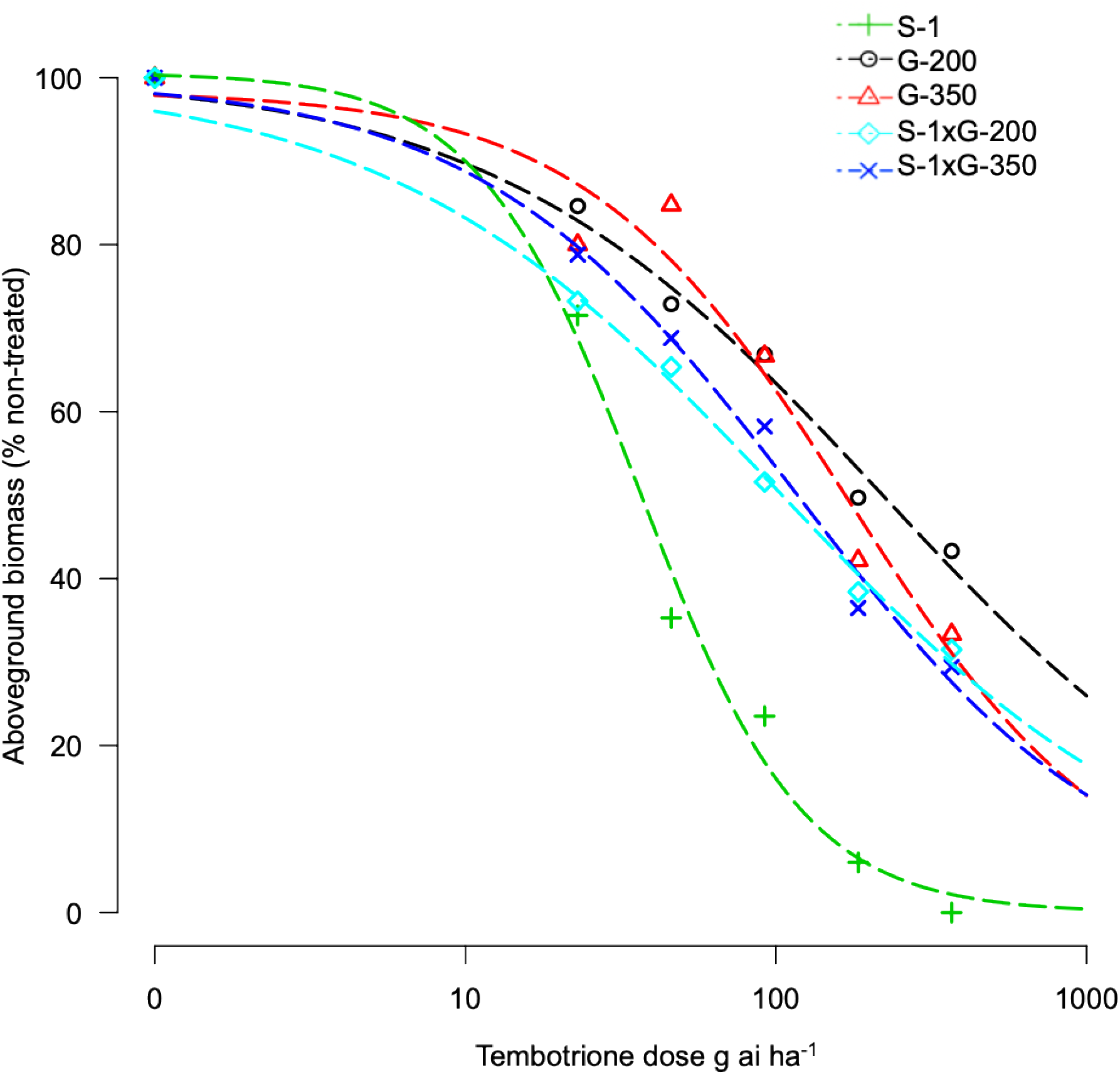
Tembotrione dose-response curves representing the above ground dry biomass of S-1 (susceptible), F_1_ (S-1 × G-200), F_1_ (S-1 × G-350) and G-200, G-350 (resistant) using the four-parameter log-logistic model.

**Figure 6:**
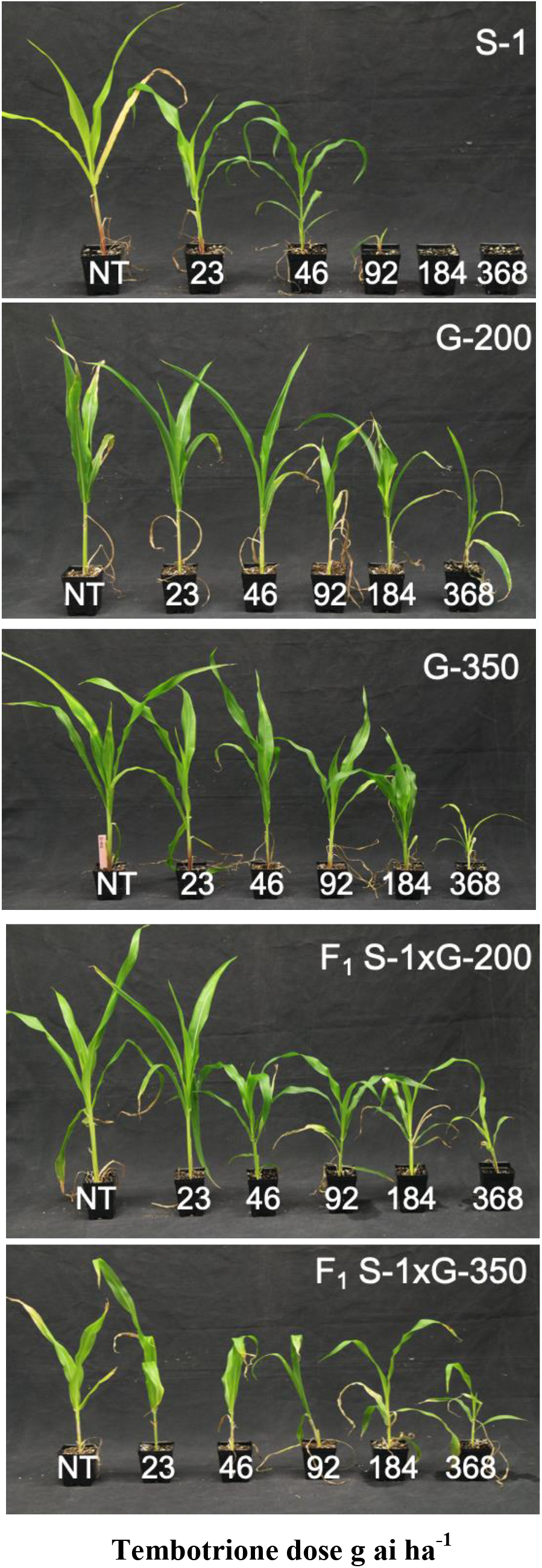
Whole plant response of parents S-1 (susceptible), G-200, G-350 (resistant), F1 (S-1 × G-200), F1 (S-1 × G-350) to different doses of tembotrione, 3 weeks after treatment. NT: Non-treated

### Mapping tembotrione resistance

To map the genomic loci controlling tembotrione resistance, a total of 208,376 SNPs were obtained using genotyping-by-sequencing from 150 F_2_ progeny (S-1 × G-200) and parents (S-1, G-200). A subset of 1,954 SNP markers polymorphic to both parents with less than 30% missing values were retained. Further, filtering for missing rate (>90%), strong segregation distortion, marker distribution and redundant markers resulted in a total of 696 markers that were used for construction of a linkage map. The map of 1021 cM was prepared which had an average distance of 1.7cM between two adjacent markers. A total of 8 QTLs on chromosomes 2, 3, 4, and 8 were mapped with a high LOD score (LOD>2) (Figure 7; Table 5) for two traits, recovery (RE) i.e difference between leaf chlorophyll content at 2 and 4 WAT, and visual scoring at 3 WAT (VI) obtained from 150 F_2_ plants (Figure S1). The LOD score of detected QTLs ranged from 2.0 to 6.0 and the phenotypic variations explained (PVE) values ranged from 9 to 44%.

**Table 5.** Quantitative trait loci (QTLs) detected for the recovery (RE) and visual injury 3 weeks after treatment (VI) along with logarithm of the odds (LOD) and phenotypic variation explained (PVE) explained by QTLs.

| Trait | QTL | Left Marker | Right Marker | LOD | PVE (%) | Add | Previously Reported Trait | Reference |
| --- | --- | --- | --- | --- | --- | --- | --- | --- |
| RE | qRE8.1 | Ch8_20217087 | Ch8_20519857 | 4.32 | 44.35 | -1.97 | Efficiency of PSII reaction centers, Chlorophyll fluorescence | Ortiz et al 2017, Fiedler et al 2016 |
| RE | qRE4.1 | Ch4_13622192 | Ch4_14717522 | 2.15 | 8.26 | 4.92 | Photochemical quenching | Ortiz et al 2017, Fiedler et al 2016 |
| RE | qRE4.2 | Ch4_33879202 | Ch4_35080912 | 2.14 | 36.56 | -1.19 | - | - |
| RE | qRE8.2 | Ch8_18330711 | Ch8_20217087 | 2.06 | 26.80 | -3.64 | Efficiency of energy captured by open PSII reaction centers, Chlorophyll fluorescence | - |
| VI | qVI4.0 | Ch4_46023941 | Ch4_48400958 | 6.20 | 21.23 | 17.18 | - | - |
| VI | qVI2.1 | Ch2_45821038 | Ch2_46847104 | 3.15 | 9.71 | 11.79 | - | - |
| VI | qVI2.2 | Ch2_45308362 | Ch2_45821038 | 2.92 | 9.49 | 9.69 | - | - |
| VI | qVI3.0 | Ch3_28379809 | Ch3_29928561 | 2.82 | 10.11 | 12.18 | Chlorophyll Content | Fiedler et al 2016 |

**Figure 7:**
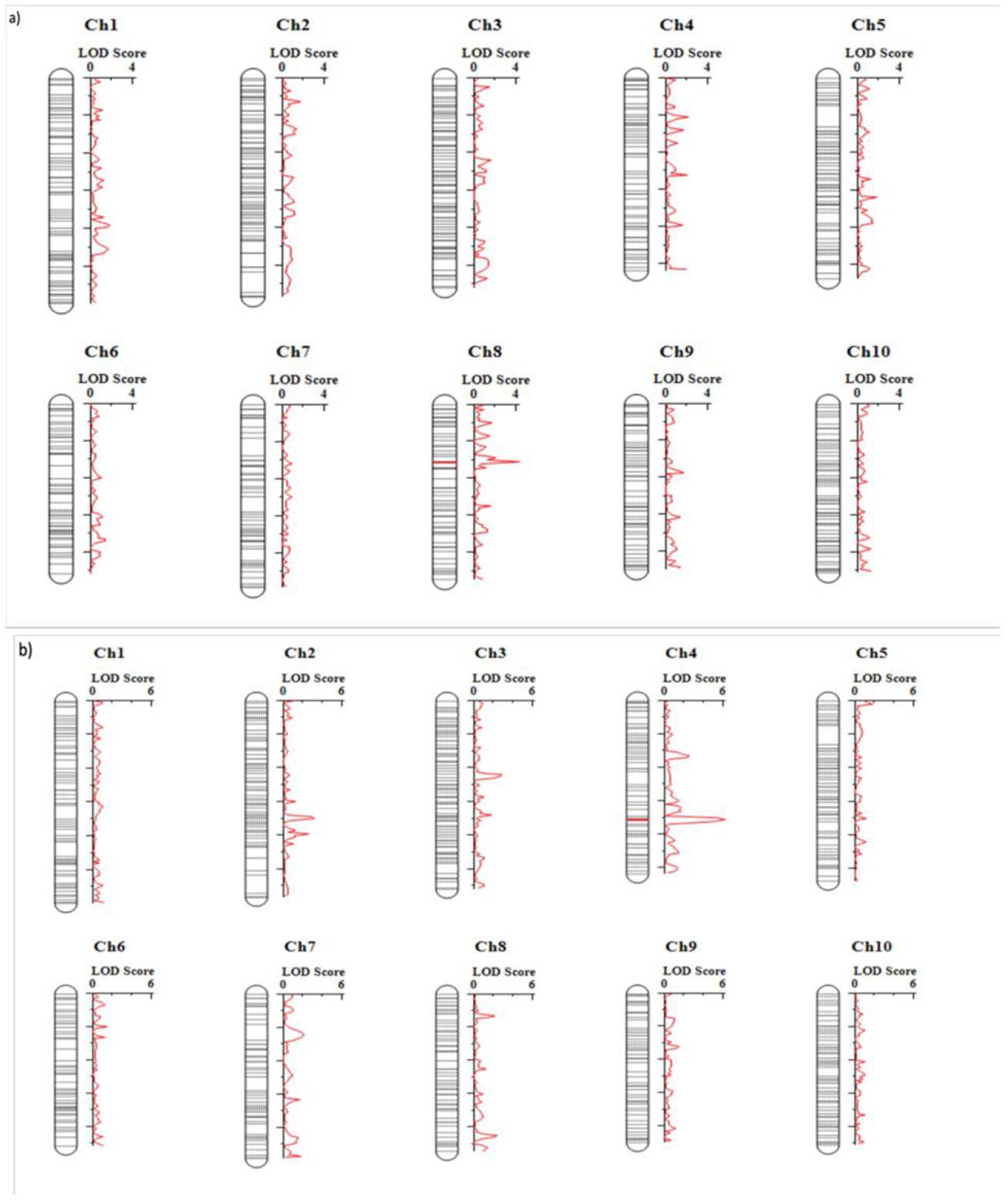
Quantitative trait loci (QTLs) detected from analysis of 150 plants from a single F_2_ family of S-1 × G-200 for different traits: a) Recovery (RE); and b) visual injury (VI) at 21 days after treatment.

## Discussion

Tembotrione, an HPPD-inhibitor, is widely used in corn for POST control of both grasses and broad-leaved weeds, but not registered for use in sorghum due to crop injury (Damalas *et al.*, 2018). In this study, we identified two sorghum genotypes, G-200 and G-350, with a high level of resistance and one genotype S-1 sensitive to tembotrione from SAP. Even though cultivated sorghum hybrids are susceptible to tembotrione; variable levels of responses was reported in sorghum germplasm upon treatment with tembotrione (Cunha *et al.*, 2016). The genotypes, G-200 and G-350 were found ~10- and 7-fold more resistant, respectively, under greenhouse and ~2-fold more resistant under field conditions as compared to S-1 and Pioneer 84G62 (Table 1). In susceptible genotypes, application of tembotrione causes foliar bleaching, leaf necrosis followed by complete plant death (Galon *et al.*, 2016). Interaction of genotypes with environmental factors such as rainfall, soil type, and weather conditions play a key role in plant response to HPPD-inhibitors (Bollman *et al.*, 2008), which may explain the difference in the level of resistance between greenhouse and field conditions. Variation in the efficacy of HPPD-inhibitors in response to environmental factors such as temperature and relative humidity were reported in several weeds (Johnson and Young, 2002; Godar *et al.*, 2015). However, in both greenhouse and field conditions, G-200 and G-350 survived the field dose of tembotrione; whereas, S-1 or Pioneer 84G62 were severely injured (Table 1).

The natural resistance to tembotrione in G-250 and G-350 appears to be not conferred by any alteration to the molecular target of this herbicide, i.e. *HPPD* gene, because no difference in *HPPD* gene sequence was found between the resistant (G-250 and G-300) or sensitive (S-1) genotype (Data S1). Likewise, no naturally evolved mutations in the *HPPD* gene that confer resistance to HPPD-inhibitors were found in plants (Lu *et al.*, 2020). Nonetheless, recently, soybean varieties resistant to HPPD-inhibitors were developed through transgenic technology by inserting an insensitive *HPPD* gene (Siehl *et al.*, 2014; Dreesen *et al.*, 2018).

The CYP enzymes are known to metabolize HPPD-inhibitors, such as mesotrione (Ma *et al.*, 2013; Nakka *et al.*, 2017), tembotrione (Küpper *et al.*, 2018), or topramezone (Elmore *et al.*, 2015) in naturally evolved resistant weed biotypes. In this research, the genotypes, G-200, and G-350 exhibited significant biomass reduction in response to pretreatment with CYP-inhibitors, malathion, or PBO followed by tembotrione application, suggesting that tembotrione is metabolized by CYP activity (Figures 3 and 4). Similarly, use of these inhibitors followed by tembotrione, showed ~10% more biomass reduction compared to tembotrione alone treatments in a tembotrione-resistant common waterhemp population (Oliveira *et al.*, 2018). Metabolism of tembotrione by hydroxylation followed by glycosylation, catalyzed by CYPs has also been reported in a tembotrione-resistant Palmer amaranth biotype (Küpper *et al.*, 2018). Furthermore, RNA-Seq analysis revealed differential expression of several CYP genes, for example, 3-4 fold upregulation of *CYP72A219, CYP81E8*, respectively, was found in the same above tembotrione-resistant Palmer amaranth biotype (Küpper, 2018). In corn, multiple CYPs located in *nsf1* locus were found to metabolize tembotrione, mesotrione, as well as other herbicides (Williams and Pataky, 2010).

Based on the response of F_1_ progeny (S-1 × G-200 and S-1 × G-350) to tembotrione treatment, we found that the tembotrione resistance in G-200 and G-350 is a semi-dominant trait (Figures 5 and 6; Table 2). Furthermore, F_2_ data demonstrated that this resistance is controlled by multiple genes (Table 4). Genetic analyses of sweet corn inbred lines revealed a single recessive allele controlling the sensitivity to tembotrione (Williams and Pataky, 2008). The genetic basis of tembotrione resistance is not extensively studied in plants; however, mesotrione (another widely used HPPD-inhibitor) resistance in several common waterhemp populations across US Midwest (Huffman *et al.*, 2015; Kohlhase *et al.*, 2018; Olivera *et al.*, 2018) was found to be inherited by a semi-dominant polygenic trait.

We mapped eight QTLs associated with tembotrione resistance on chromosomes 2,3,4, and 8 using the sequence data from F_2_ progeny (Figure 7). To our knowledge, this is the first report of QTLs associated with tembotrione resistance in grain sorghum. These QTLs were previously reported for other traits in sorghum related to chlorophyll fluorescence (Fiedler *et al.*, 2014), photochemical quenching (Ortiz *et al.*, 2017) (Table 5). Experiments are in progress to further fine map and identify the precise location of the gene(s) responsible for tembotrione resistance in grain sorghum.

As mentioned earlier, our data indicates that the tembotrione resistance is a polygenic trait, and such traits can express differently in different genetic backgrounds. Therefore, tembotrione resistance can potentially be improved by crossing G-200 and G-350 or with other commercial genetic backgrounds. Such work has been reported to enhance performance of quantitative traits in different genetic backgrounds and environmental conditions such as drought (Reddy *et al.*, 2009), stay green (Subudhi *et al.*, 2000), cold tolerance (Knoll and Ejeta, 2008), and yield (Nagaraja Reddy *et al.*, 2013) in grain sorghum. Therefore, there is enormous potential for improving tembotrione resistance by testing the expression of this trait in different genetic backgrounds and for the development of tembotrione-resistant sorghum varieties.

Since sorghum can outcross with closely-related wild and weedy species, such as johnsongrass or shattercane, one of the major concerns of development of herbicide-resistant sorghum varieties has been natural transfer of such resistance into these weed species (Ohadi *et al.*, 2017). However, recent reports suggest that the outcrossing rate of sorghum with johnsongrass was as low as ~1% under controlled conditions (Hodnett *et al.*, 2019) and 2-16% with shattercane under field conditions (Schmidt *et al.*, 2013). Although the possibility of outcrossing is minimal, if an herbicide resistance trait escapes into the wild species, necessary stewardship practices must be developed and integrated into sorghum weed management practices.

## Conclusions

In conclusion, we have identified sorghum genotypes (G-200 and G-350) with natural resistance to tembotrione from the sorghum association panel, which can potentially be used to introgress the tembotrione resistance into breeding lines by conventional or marker-assisted breeding methods. CYP-inhibitor assay suggested CYP-mediated metabolism of tembotrione in the resistant genotypes. Genetic analyses of F1 and F2 progeny demonstrated that the resistance is a semi-dominant polygenic trait. Furthermore, GBS-based QTL mapping revealed eight QTLs associated with tembotrione resistance in grain sorghum. Future research needs to be focused on incorporating the resistant trait with elite breeding varieties, testing the hybrid performance, and improving herbicide resistance in high yielding and stress tolerance hybrids.

## Materials and Methods

### Plant materials

Sorghum genotypes from the SAP (Casa *et al.*, 2008) were used in this study. A commercial sorghum hybrid Pioneer 84G62, and a corn inbred B73 (naturally resistant to tembotrione) were also used for comparison.

### *In vitro* screening

Sorghum genotypes (~317) from SAP along with Pioneer 84G62 and B73 were used for initial screening with tembotrione under *in vitro* conditions. Seeds of the all genotypes were germinated in plastic Petri dishes (100 diameter × 20 mm height) containing 0.8% w/v solidified agar medium (PhytoTech Laboratories, Lexana, Kansas, USA). Seeds were surface sterilized with 2% ethanol for 2 min followed by 15% bleach for 15 min. Subsequently, seeds were rinsed 2 to 3 times with sterile distilled water before placing them on the agar medium. About 8 to 10 seeds were placed in each Petri dish for germination and incubated in a growth chamber maintained at 24℃ with 16 h day/night photoperiod under a photosynthetic flux of 200 μmol m^−2^ s^−1^ (daylight fluorescent tubes). Upon germination, seedlings at 1 to 2 leaf stage were transferred to culture vessels (PhytoTech Laboratories, Lenexa, Kansas, USA) containing solidified agar supplemented with 0.25 μM molecular grade tembotrione. All transplanted culture vessels were incubated in the same growth chamber, maintained at the same conditions as indicated above. The experiment was conducted once with 2 replications (2 to 4 plants in each culture vessel). The response of genotypes to tembotrione treatment was evaluated visually (percent injury) at 2 and 4 WAT based on a 0 to 100% rating scale (0% is no injury and 100% is complete death) (Abit *et al.*, 2009).

### Whole plant assay

Ten sorghum genotypes (Table S1) that exhibited minimum tembotrione injury under *in vitro* conditions were tested for their response to tembotrione under greenhouse conditions. The seeds of sorghum genotypes were planted in square pots (15 × 15 × 15 cm) filled with a potting mixture (ProMix Ultimate, Premier Tech Horticulture, Mississauga, Ontario, Canada). The seedlings at 2-3 leaf stage (Roozeboom and Prasad, 2019), were transplanted in square pots (6 × 6 × 6 cm) and grown in a greenhouse maintained at 25/20℃, 15/9 hours day/night photoperiod with a hotosynthetic photon flux density of 750 μmol m^−2^ s^−1^ and relative humidity of 60 ± 10 percent. The plants were fertilized (Miracle GRO® All-purpose plant food, ScottsMiracle-Gro, Marysville, Ohio, USA) as needed. The sorghum seedlings at 4-5 leaf-stage (Roozeboom and Prasad, 2019) were treated with tembotrione (LAUDIS^®^, Bayer Crop Science, USA; https://www.cropscience.bayer.com) at 92 g ai ha^−1^ with 0.25% methylated soy oil (DESTINY^®1^, WinField^®1^United, https://www.winfieldunited.com/) using a bench-top track spray chamber (Generation III, De Vries Manufacturing, Hollandale, Minnesota, USA) equipped with a single flat-fan nozzle (80015LP TeeJet tip, Spraying Systems Co., Wheaton, Illinois, USA) delivering 187 L ha^−1^. Each plant was considered as an experimental unit, eight replications were used for each genotype. The response of sorghum genotypes to tembotrione treatment was evaluated by visual injury rating as described above (Abit *et al.*, 2009). The above-ground plant biomass was harvested 3 WAT and dried in an oven at 60°C for 72 h. The weight of dried biomass was recorded as described later in a separate section. The experiment was repeated two times following same procedure and growth conditions.

### Dose-response assay

Two genotypes i.e., G-200 and G-350 that exhibited the least injury, and one highly sensitive genotype S-1 that exhibited the highest injury to tembotrione treatment identified from *in vitro* and whole plant assays, along with Pioneer 84G62 were tested in a tembotrione dose-response study to determine the level of resistance. The sorghum genotypes were treated with tembotrione at 0, 5.75, 11.5, 23, 46, 92, 184, 368, 736 g ai ha^−1^. The experiment was conducted following the same plant growth conditions and herbicide application procedure as described above in the whole plant assay. The experiment was conducted in a completely randomized design (CRD) with four replications and repeated twice. The aboveground plant biomass reduction was measured as described above.

### Field testing

Upon confirmation of the level of resistance to tembotrione in the greenhouse, the tembotrione-resistant sorghum genotypes G-200 and G-350 were evaluated under field conditions. Experiments were conducted in summer of 2017 at two KSU research sites; Ashland Bottoms Research Farm, Manhattan (Reading silt loam soil type; Pachic Agriustolls taxonomic class), and Agricultural Research Center, Hays (Harney silt loam soil type; Typic Agriustolls taxonomic class). *S*-metolachlor at 2 kg ai ha^−1^ was applied as a pre-emergence herbicide to all plots at both sites to suppress existing weeds in the field before planting sorghum. Seeds of sorghum genotypes were planted in both locations on June 6, 2017 with a 76 cm space between rows and 7.6 cm space between plants, and 2.5 cm planting depth. The experimental plots were 3 m wide and 6 m long with four rows; the resistant or susceptible genotypes were planted in the middle two rows along with two border rows planted with Pioneer 84G62 to avoid herbicide drift from nearby treatments. POST application of tembotrione was made to individual plots when the sorghum plants reached 3 to 5 leaf collar stages (Roozeboom and Prasad, 2019). Tembotrione treatments included 0, 92, 184, and 368 g ai ha^−1^. Herbicides were applied using a CO_2_-powered backpack-type research sprayer equipped with TurboTee 11002 nozzles calibrated to deliver 140 l ha^−1^ at 234 KPa. Experiments were conducted in a randomized complete block design with factorial arrangement with sorghum genotype and herbicide dose as the two factors. All treatments were replicated four times at each site. Sorghum response to herbicide treatments was visually assessed 1, 2, 4, and 8 WAT using a scale of 0 (no visible injury) to 100% (plant death) compared to the non-treated plants. At physiological maturity, grain weight was measured for each genotype. Two sorghum heads were collected for each genotype separately from all treatments and replications and dried in an oven at 60°C for 1 week. The dried sorghum heads from each plant were subsequently threshed to determine grain yield from a single plant.

### Generation and evaluation of F1 and F2 progeny

To study the inheritance and mapping of tembotrione resistance, direct and reciprocal crosses were performed using tembotrione-resistant (G-200 and G-350) and - sensitive (S-1) genotypes in a crossing nursery at KSU research farm, Ashland Bottoms, KS. The crosses were made using the plastic bag method (Rakshit and Bellundagi, 2019). The F_1_ seeds were harvested from individual plants. The F_1_ progeny from S-1 × G-200 and S-1 xG-350 were evaluated in a tembotrione dose-response assay by treating the plants with 0, 23, 46, 92, 184, 368 g ai ha^−1^ of tembotrione. Each plant was considered as an experimental unit with 10 to 12 replications per dose. The same procedure as described above was followed for tembotrione dose-response assay of F_1_ progeny. Three F_1_ plants per cross that exhibited resistance to tembotrione, were selected to generate F_2_ seeds by self-pollination.

The F_2_ progeny were evaluated under greenhouse conditions with a single dose of tembotrione to determine the segregation of resistant and susceptible plants. Approximately 150 seedlings from a single F_2_ family (up to two F_2_ families) along with the parents were raised in the greenhouse (as described above under the same growth conditions). The seedlings (4-5 leaf stage) were treated with 276 g ai ha^−1^ of tembotrione following the same procedure as described above. The response of F_2_ plants was assessed by visual injury rating (as described above) at two and three WAT (Abit *et al.*, 2009). Further, plants were grouped as highly injured/dead (susceptible) or minor/no symptoms (resistant) at four WAT in comparison with the parental genotypes. In addition, total leaf chlorophyll content was estimated in parents and F_2_ progeny on three and four WAT. Chlorophyll content was measured at three different spots on the leaf blade along the length of the youngest fully opened leaf using a self-calibrating soil plant analysis development (SPAD) chlorophyll meter (Konica Minolta SPAD 502 Chlorophyll Meter, Chiyoda City, Tokyo, Japan). The chlorophyll content obtained from the three spots were averaged and used as total leaf chlorophyll content. However, the leaf chlorophyll index was recorded from the second run of S-1 × G-200 F_2_ evaluation which was used for the QTL mapping experiment (described later in separate section).

### *HPPD*-gene sequencing

The *HPPD* gene from G-200, G-350, and S-1 were sequenced to determine if any target site alterations confer resistance to tembotrione. Leaf tissue (3-4 leaf stage plants) was collected from three plants of each genotype grown in the greenhouse as described above and under similar growth conditions. The genomic DNA was extracted using GeneJET™ Plant Genomic DNA Purification Mini Kit (Thermo Scientific™, Waltham, Massachusetts, USA) following the manufacturer’s instructions. The concentration of the DNA samples were quantified using NanoDrop™ (Thermo Scientific™, Waltham, Massachusetts, USA). The sorghum *HPPD* gene ~2kb was amplified using the primers Sg_HPPD F (5’GACACGATGAATGCCCATGC 3’) and Sg_HPPD R (5’ AGAGAGATGACAGTACAGTGTTGT 3’) designed from Sobic.002G104200.1 in the sorghum reference genome V3.1.1 (McCormick *et al.*, 2017). Polymerase Chain reaction (PCR) was performed using T100™ Thermal Cycler (Bio-Rad Inc., Hercules, California, USA). The PCR mixture contained 50-80 ng of gDNA, 0.5μM each of forward, reverse primer, and 1x of GoTaq® G2 Green Master Mix (Promega™, Madison, Wisconsin, USA). PCR amplification was done using the following PCR cycling conditions, initial denaturation: 94 °C for 5 min, followed by 35 cycles of denaturation: 94°C for the 30s, annealing 60°C for 45s and extension: 72°C for 45s and final extension: 72°C 7 mins. The PCR products were analyzed in 1.5% agarose gel to confirm the targeted amplicon size and purified using GeneJET™ PCR Purification Kit (Thermo scientific™, Waltham, Massachusetts, USA) The PCR purified samples were sequenced by Sanger sequencing service provided by GENEWIZ, LLC., New Jersey, South Plainfield, USA. The sequences were aligned using Clustal Omega multiple sequence alignment tool (EMBL-EBI) to check for the mutations.

### CYP-inhibitor study

To determine if CYP-mediated metabolism of tembotrione confers resistance in G-200 and G-350 genotypes, experiments were conducted using two CYP-inhibitors, malathion and PBO. The sorghum genotypes G-200, G-350, S-1 along with Pioneer 84G62 and a corn genotype B73, were grown in the greenhouse (as described above and under similar growth conditions). Malathion (SPECTRACIDE® MALATHION INSECT SPRAY CONCENTRATE, Spectrum Brands, Inc, https://www.spectracide.com/) at 0, 2000, and 4000 g ai ha^−1^ or PBO (Fisher Scientific, Waltham, Massachusetts, USA) at 4500 g ai ha^−1^ along with 0.25% non-ionic surfactant (NIS) was applied one hour prior to tembotrione treatment. Soil drenching of 5mM malathion 24 hours after primary application as a booster dose was given only for the malathion treatments. Tembotrione was applied at 0, 92, 184, and 368 g ai ha ^−1^. All the treatments were arranged in a factorial design. The same procedure as mentioned in the above tembotrione dose-response assay was followed for chemical treatments (malathion, PBO, and tembotrione) and data collection.

### Genotyping by sequencing

Approximately 150 plants of a single F_2_ family derived from S-1 × G-200 along with parents, were grown in the greenhouse as described above and under the same growth conditions. An equal amount (two 2cm leaf bits; ~150 mg) of leaf tissue was collected from all plants in 96-deep well plates. One 3.2-mm stainless steel bead was added to each well and the leaf tissue was ground for 3 min at 20 cycles per sec to obtain fine powder in a Mixer Mill (Retsch GmbH, Haan, North Rhine-Westphalia, Germany). Genomic DNA was extracted using the Cetyltrimethylammonium bromide (CTAB) method (Bai *et al.*, 1999) with minor modifications. The DNA concentration in the extracted samples was quantified by FLUOstar Omega microplate reader (BMG LABTECH, Ortenberg, Baden-Württemberg, Germany) using a Quant-iT™ PicoGreen® dsDNA Assay Kit (Life Technologies, Grand Island, New York, USA). Each sample was normalized to contain 10 ng/μl DNA using QIAgility Liquid Handling System (Qiagen, Germantown, Maryland, USA) for library construction. Approximately 150 ng of genomic DNA of each sample was used to construct a library following the tGBS^®^ protocol (Ott *et al.*, 2017) with modifications, and the DNA library was sequenced on the HiseqX 10 platform at Novogene Corporation Inc., Sacramento, California, USA. Sequencing reads were trimmed and de-barcoded using the pipeline described in the previous tGBS^®^ study (Ott *et al.*, 2017). Clean reads of each sample were aligned to *Sorghum bicolor* genome (Genbank accession GCA_000003195.3) (Paterson *et al.*, 2009) using BWA v0.7.12-r1039 (Li and Durbin, 2009) and unique mapped reads were retained for variant discovery using HaplotypeCaller in GATK v4.1(McKenna *et al.*, 2010). GATK SelectVariants with parameters (-select-type SNP --restrict-alleles-to BIALLELIC -select ‘QD ≥ 10.0’ -select ‘DP ≥ 200.0’) was applied to filter variants. The SNPs were converted to ABH format (“A” represents resistant parent allele, “B” represents susceptible allele and “H” represents heterozygous allele) and only polymorphic SNPs between the R and S genotypes were retained using a custom-made Microsoft Excel template. The filtered polymorphic SNPs were used for the construction of a linkage map and QTL analysis.

### Linkage and QTL mapping

The linkage map was obtained using QTL IciMapping (version 4.5). The grouping and ordering of 606 polymorphic SNP markers were carried out using the regression mapping algorithm RECORD (REcombination Counting and ORDering) based on recombination events between adjacent markers. Further, rippling was done for fine-tuning of the ordered markers on their respective chromosomes by the sum of adjacent recombination fractions (SARF) algorithm with a default window size. The QTL mapping for RE and VI was performed using the inclusive composite interval mapping (ICIM) method with additive assumption was performed using the QTL IciMapping (version 4.5) (Meng *et al.*, 2015). The LOD significance thresholds (*P* < 0.05) were determined by running 1000 permutations (Churchill and Doerge, 1994). Previously reported QTLs for similar regions were obtained from sorghum QTL Atlas (Mace *et al.*, 2019). The QTLs (q) were named based on the trait abbreviation followed by the chromosome number.

### Statistical analysis

Dry biomass (% of non-treated) was calculated following the formula:

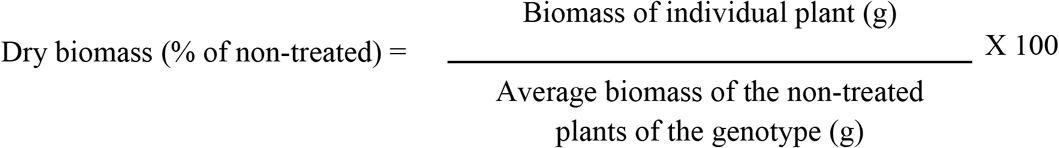

Tembotrione dose-response data expressed as dry biomass (% of non-treated) or percent injury were subjected to non-linear regression analysis using a three or four-parameter log-logistic model using a ‘drc’ (Ritz *et al.*, 2005) package in R (Development Core Team, 2013) following (Knezevic *et al.*, 2007; Shyam *et al.*, 2019) to estimate GR_50_ or ID_50_. A “Lack-of-fit” test was performed using the “model fit” function of ‘drc’ to assess the fit of data to various regression models. Differences between the estimated GR_50_ or ID_50_ values were tested with each other by t-test using the “compParm” function in the ‘drc’ package. The dose-response curves were generated using the ‘plot’ function in the ‘drc’ package.

Analysis of variance (ANOVA) was performed following Fisher’s LSD test to separate means and significance at p ≤ 0.05 using the ‘agricole’ package in R (de Mendiburu, 2014). The plots were generated using the ‘R’ package ‘ggplot2’ (Wickham and Wickham, 2007). A Chi-square (χ2) goodness of fit test (Cochran, 1952) was used to fit to a single dominant gene by comparing the observed and expected segregation frequencies of tembotrione resistant or - susceptible plants.

## Supplemental Material

Additional supporting information.

**Supplemental Table S1.** Sorghum genotypes that exhibited minimal visual injury in the *in vitro* screening.

**Supplemental Table S2.** Response of sorghum genotypes to whole plant assay.

**Supplemental Data S1.** Sequence alignment of *HPPD-*gene.

**Supplemental Figure S1.** Phenotypic Distribution of S-1x G-350 F_2_ progeny and parents for a) Recovery (RE); and b) Visual injury at four weeks after treatment (VI).

**Supplemental Figure S2.** Linkage map obtained from 606 markers spread across the sorghum genome.

